# Revealing within-species diversity in uncultured human gut bacteria with single-cell long-read sequencing

**DOI:** 10.1101/2022.03.24.485581

**Authors:** Masato Kogawa, Yohei Nishikawa, Tatsuya Saeki, Takuya Yoda, Koji Arikawa, Haruko Takeyama, Masahito Hosokawa

**Affiliations:** Research Organization for Nano and Life Innovation, Waseda University, 513 Wasedatsurumaki-cho, Shinjuku-ku, Tokyo 162-0041, Japan; Computational Bio Big-Data Open Innovation Laboratory, National Institute of Advanced Industrial Science and Technology, 3-4-1 Okubo, Shinjuku-ku, Tokyo 169-8555, Japan; bitBiome, Inc., 513 Wasedatsurumaki-cho, Shinjuku-ku, Tokyo 162-0041, Japan; Department of Life Science and Medical Bioscience, Waseda University, 2-2 Wakamatsu-cho, Shinjuku-ku, Tokyo 162-8480, Japan; Institute for Advanced Research of Biosystem Dynamics, Waseda Research Institute for Science and Engineering, 3-4-1 Okubo, Shinjuku-ku, Tokyo 169-8555, Japan

## Abstract

Bacterial genome structure changes dynamically, and structural variants can change bacterial phenotype; However, obtaining the complete genome and analyzing genome structure of uncultured bacteria has been challenging. We aimed to develop a single-cell amplified genome long-read assembly (scALA) workflow to construct circular single-cell amplified genomes (cSAGs) from long-read single-cell sequencing data of targeted uncultured bacteria. In particular, scALA generated cSAGs from nanopore long-read sequencing data of SAGs by producing contiguous sequences with repeated bias reduction and assembly processes. From 12 human fecal samples, scALA generated 16 cSAGs of three specifically targeted bacterial species, *Anaerostipes hadrus, Agathobacter rectalis*, and *Ruminococcus gnavus. A. hadrus* cSAGs exhibited large, ten kbp-long, phage insertions, saccharide metabolic capacity, and frequent genomic recombination with related strains from cohabitant hosts. Noteworthy, cSAGs constructed using this method could expand bacterial genome databases and our understanding of within-species diversities in uncultured bacteria.

## Introduction

Culture-independent bacterial genome analysis has revealed that gut microbiota plays a crucial role in controlling host physiology and metabolism. The analysis relies on the availability of reference bacterial genomes for accurate functional assignments of specific organisms and taxonomic classification of microbiota. However, an estimated half of human gut microbiota species lack reference genomes. In recent years, short-read sequencing with assembly and binning algorithms has led to an explosion of metagenomic assembled genomes (MAGs), with more than 100,000 MAGs created in individual studies^1–3^. However, because MAG bins represent population consensus genomes, most have low assembly quality, including unlinked loci, missing rRNA genes, and chimeric sequences^4–6^. This genome incompleteness has raised concerns about the quality of MAG-derived reference databases and the validity of MAG-based studies. In particular, gut bacterial species remain poorly characterized in terms of intra-species diversity. Thus, a novel genome recovery approach is required to enhance the reliability of reference genomes and unveil within-species variations in yet uncultured bacteria.

Single-cell genomics is an alternative approach for culture-independent recovery of bacterial genomes^7^. In contrast to metagenomics, single-cell genomics does not require bacterial population clonality because it recovers genome sequences from individual cells. In this process, a single bacterial cell is isolated, lysed, and the entire genome is amplified. Moreover, multiple displacement amplification (MDA)^8^ generates sufficient replicating DNA with high fidelity and large fragment sizes; however, certain problems arise. For example, MDA typically introduces chimeric artifacts within a single genome by linking non-contiguous genomic regions^9^. A substantial bias in genome coverage has also been observed, leading to a lack of coverage of certain genomic regions. Consequently, single-cell amplified genomes (SAGs) often have fragmented and incomplete sequences containing errors and can be misinterpreted similarly to MAGs. Under the Minimum Information about a Single Amplified Genome standards^5^, only a minimal number of high-quality draft SAGs and circular SAGs (cSAGs) have been recovered to date^10^.

In this study, we used high-throughput single-cell genome sequencing to recover complete genomes from uncultured human gut microbiota. We combined a massively parallel single bacterial genome sequencing technique, namely single-cell genomes amplified using gel beads sequencing (SAG-gel)^11^ platform, with a nanopore long-read sequencer^12,13^, and investigated three uncultured human gut bacterial species of the order Clostridiales.

## Results

### Evaluation of the conventional long-lead assembly pipeline using *Escherichia coli* single-cell genomes

Initially, we evaluated long-read SAGs (LR-SAGs) constructed from *E. coli* single-cell genomes using existing long-read assemblers to identify challenges in *de novo* assembly of single-cell long-read sequences (scLRs; Table S1).

First, 96 SAGs of *E. coli* K12 were prepared using SAG-gel platform^11^, and five amplified genomes were then randomly selected and individually sequenced using a nanopore sequencer. ScLRs (300 Mb each) were merged into a single file. LR-SAG obtained from scLRs using Flye^14^, one of the most reliable long-read assemblers for sequencing genomes extracted from cultured bacteria, had a genome completeness of 1.53 % and lost most sequence information. Similarly, Miniasm^15^ output a low-quality LR-SAG with a genome completeness of 1.17 %. In contrast, Canu^16^, one of the earliest long-read assemblers which has a built-in chimeric sequence processing unit, constructed LR-SAG with a high genome completeness of 87.5 % compared with that constructed using the previous two assemblers. However, the number of contigs was 61, including numerous fragments with the same sequence information, as indicated by the duplication ratio of 1.039. Therefore, we first removed chimeras from LR-SAGs using Canu and then assembled them using Flye and Miniasm, resulting in a reduced number of contigs (39 and 47, respectively), while maintaining genome completeness of > 70 %. However, both assembly results showed a maximum genome completeness of 87.5 %, confirming that a substantial amount of scLR sequence information was lost. Because mostly scLR sequence regions with small relative depth in LR-SAGs exhibited missing sequence information (Fig. S1), we hypothesized that improving sequencing biases in scLRs would enhance the assembly quality.

### Enhancing genome completeness by reducing bias in scLRs

We developed single-cell amplified genome long-read assembly (scALA) pipeline to assemble bacterial draft genomes from scLRs with amplification bias and chimeric sequences (Fig. 1). This pipeline first removed low-quality reads from the input scLRs and then constructed reference contigs as intermediate reference LR-SAGs for the sequence debiasing process. Following chimera removal using Canu, scALA assembled input scLRs to intermediate reference contigs using Flye. The input scLRs were then mapped against the reference contigs to identify the area mapped with considerable designated read depth, and the excess reads were removed to debias the input reads (Fig. 1A). Thereafter, the debiased LR-SAGs that improved genome coverage breadth were constructed by assembling the bias-reduced scLRs and used as renewal reference contigs for repeated debiasing and re-assembly of scLRs. The repeated cycles of debiasing and re-assembly provided contiguous sequences with improved genome completeness. Consensus sequences were then constructed from intermediate draft LR-SAGs obtained during the debiasing and assembly cycle. The final LR-SAG was obtained by polishing the consensus sequences with single-cell short-read sequences (scSR) obtained from the same sample.

**Figure 1:**
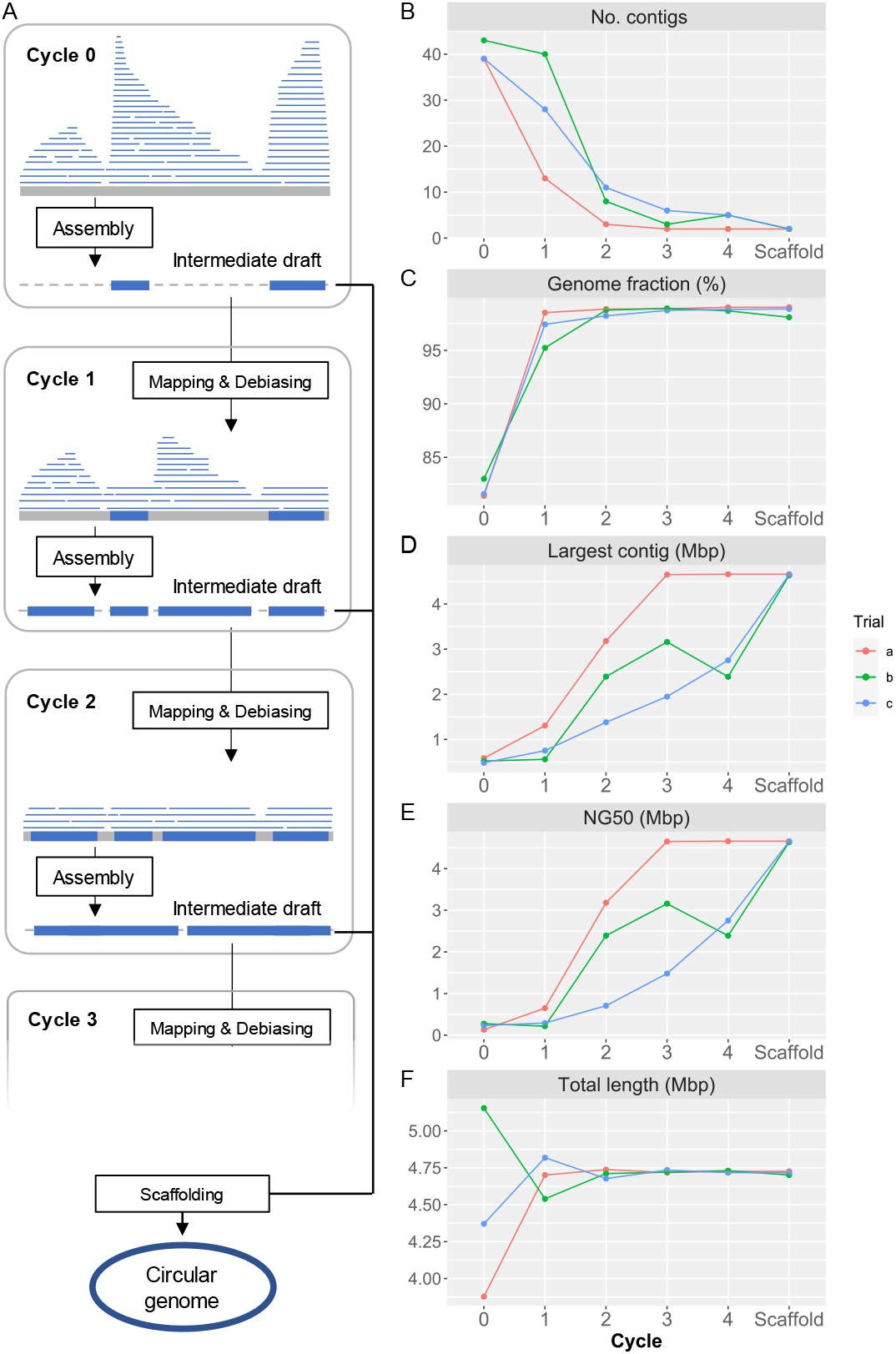
Workflow of single-cell amplified genome long-read assembly (scALA) pipeline to obtain circular bacterial genomes. **A)** The assembly of original single-cell long-read sequences (scLRs) obtained from SAGs produces a low-completeness draft genome (LR-SAG) owing to biased sequence depth caused by uneven genome amplification (Cycle 0). To improve the quality of LR-SAG, the assembly of debiased long reads is repeated by read mapping to the pre-assembled intermediate draft LR-SAG (Cycle 1, Cycle 2, …). Finally, circular LR-SAG (cSAG) is obtained by scaffolding multiple intermediate draft LR-SAGs in each cycle. **B-F)** Quality of LR-SAGs in scALA process (B: number of contigs, C: genome fraction, D: length of the largest contig, E: NG50, F: total length). The line plot color indicates each scALA analysis trial using the same *E. coli* scLR dataset.

The impact of bias reduction in scLRs on the completeness of LR-SAGs was evaluated using merged *E. coli* scLRs, which were also used to assess the conventional long-read assembly tools (Fig. 1A). After *de novo* assembly, an LR-SAG comprising 39 contigs was obtained from scLRs before debiasing with a genome completeness of 81.7 %. In contrast, an LR-SAG comprising 13 contigs was constructed from scLRs after the first debiasing with a genome completeness of 99.3 %, indicating that the bias reduction improved the genome completeness. Furthermore, the number of contigs was reduced to two after repeated debiasing, and a full-length *E. coli* genome sequence with a maximum contig length of 4.66 Mbp was finally obtained. The other contig was the F plasmid sequence from *E. coli*. These results indicate that repeated debiasing processes could construct a complete SAG from scLRs by increasing completeness and filing sequence gaps, which resulted in contig reduction.

In addition, different LR-SAGs were constructed in our pipeline for each trial (Fig. 1B-F). Three validation experiments using the same *E. coli* scLRs yielded similar LR-SAGs containing 39–43 contigs before the debiasing cycle with a genome completeness of 81.7–83.2 %. However, after four debiasing and assembly cycles, only one of the three trials resulted in constructing cSAG. This difference is because some parts of the Flye assembly algorithm use random values. Typically, random seeds are set for genome assembly to achieve reproducible assembly. However, this validation suggested that inadvertent random seed setting might prevent the acquisition of cSAGs. The alignment of LR-SAGs obtained at each step showed that sequence fragment ends were located in different genomic positions for each genome, and BLAST homology searches for each draft genome showed that the sequence fragments could be stitched together and extended as the consensus sequence (Fig. S2). Therefore, we implemented the scaffolding of all contigs obtained in different assembly cycles to generate the consensus contig and constructed contiguous cSAGs from all trials (Fig. 1B, Table S2).

### Genome comparison analysis of gut bacterial SAGs obtained from different hosts

We obtained cSAGs of three gut bacterial species obtained from 12 human fecal samples, including two cohabitant groups listed in Table S3. First, 96 short-read (SR)-SAGs of human gut bacteria were prepared using SAG gel platform^11^, and scSR data were then obtained using the HiSeq sequencing system. After *de novo* assembly of scSR, composite SR-SAGs (CSR-SAGs) were constructed by integrating SR-SAGs of the same strain^9^. Subsequently, we identified certain bacterial species whose CSR-SAG showed > 90 % completeness from multiple specimens. As shown in Fig. 2A, we targeted three bacterial species related to human health, *Anaerostipes hadrus*, *Agathobacter rectalis*, and *Ruminococcus gnavus*, which were shared among hosts^17–19^. Most CSR-SAGs of the selected species were qualified as high quality (HQ) or medium quality (MQ) and had high genome completeness (> 95 %), comprising over 100 contigs (Table S4).

**Figure 2:**
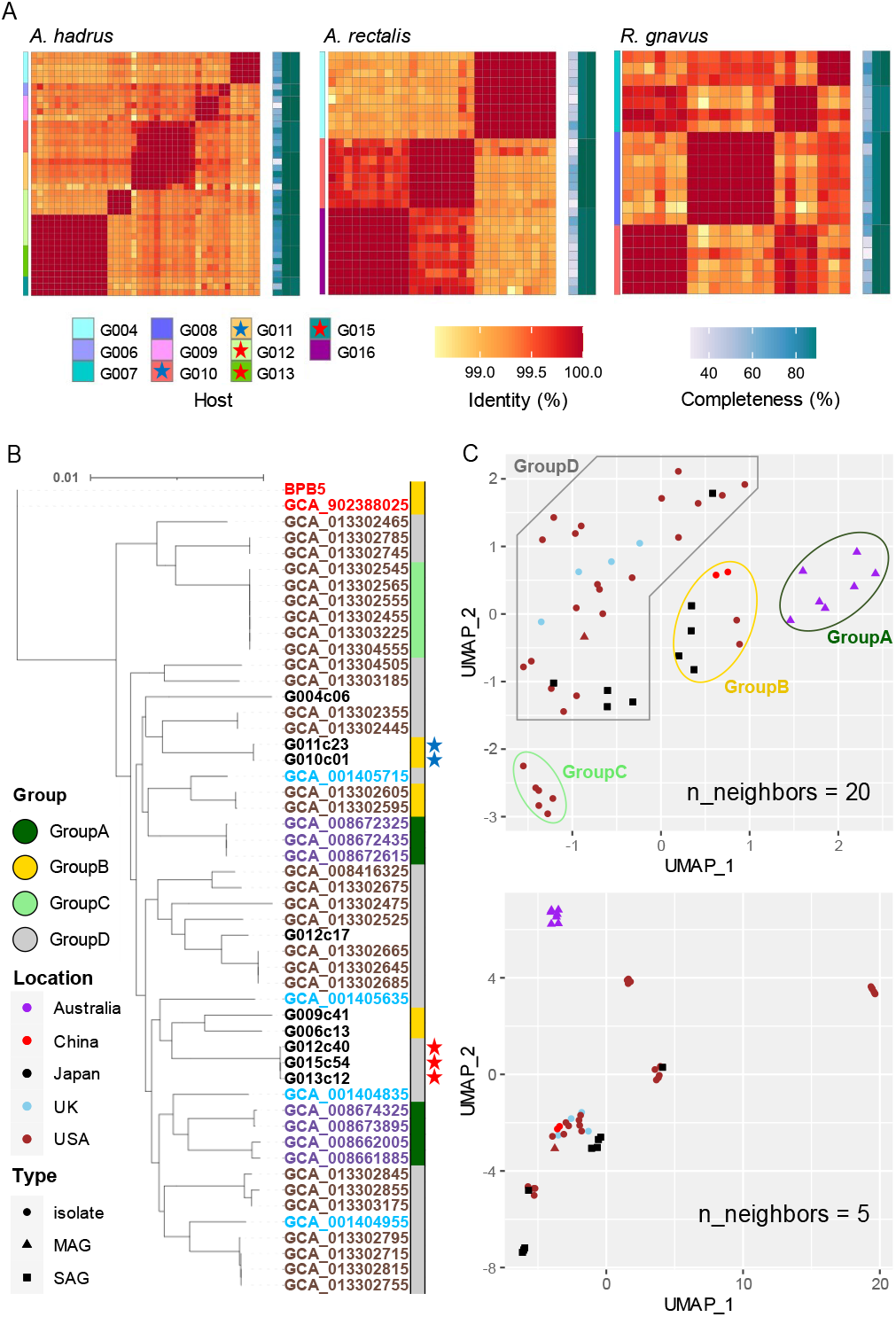
Phylogenetic analysis of targeted gut bacterial SAGs for *A. hadrus*, *A. rectalis*, and *R. gnavus*. **A)** Heatmaps of average nucleotide identity (ANI) among SAGs of the three species. Tiles on the left indicate the host of obtained SAGs. The colored stars in the host legends indicate cohabitants. Central tiles indicate the ANI between short-read (SR)-SAGs. Tiles on the right represent the completeness of SAGs processed in different steps [left column: raw SR-SAG, middle column: composite short-read SAG (CSR-SAG), right column: composite long-read SAG (LR-SAG)]. As shown in **B** and **C**, publicly available *A. hadrus* genomes obtained from isolates or metagenome-assembled genome (MAG) were compared with our *A. hadrus* SAGs. **B)** A parSNP tree of *A. hadrus* genomes based on single nucleotide polymorphisms (SNPs) of the core gene. The label color indicates the country in which the *A. hadrus* genome was obtained. Color strip on the right represents the genome group based on Uniform Manifold Approximation and Projection (UMAP) analysis (**C**). The colored stars are closely related SAGs obtained from the cohabitants. **C)** UMAP plot of *A. hadrus* genomes based on the presence of homologous gene groups. The point color indicates the country in which the bacterial sample was obtained, and point shape indicates the data type of genome sequences.

SR-SAG pairwise average nucleotide identity (ANI) clusters suggested the presence of *A. hadrus* strains shared among different hosts, the strain G010c01 and G011c23 or the strain G012c40, G013c12, and G015c54 (Fig. 2A, Fig. S3). Host groups harboring these specific strains were cohabitants. Further, we performed phylogenetic analysis within the species using *A. hadrus* comparative genome set which consisted of nine obtained CSR-SAGs and 43 *A. hadrus* draft genomes from the National Center for Biotechnology Information (NCBI) genome database. We analyzed phylogenies of these strains based on single nucleotide polymorphisms (SNPs) of the common core gene of all genomes in the dataset (Fig. 2B). The results suggested that the strain groups with > 99.5 % ANI from cohabitants were considered shared *A. hadrus* strains between couples or parents and children.

We clustered each strain genome based on the presence or absence of homologous functional genes to compare genomes from different countries. Furthermore, we used Uniform Manifold Approximation and Projection (UMAP) clustering based on Jaccard distances between genomes to classify 52 genomes into four groups (Fig. 2C, n_neighbors = 20). The *A. hadrus* genome obtained in Australia showed regionally specific clusters in UMAP analysis (Fig. 2C); however, Australian strains were distributed across multiple clades in the parSNP tree based on core gene sequences (Fig. 2B). *A. hadrus* strains G006c13, G009c41, G010c01, and G011c23, whose genomes were obtained in this research, were also located in the same UMAP cluster but different clades. Phylogenetic analysis based on core gene sequence identity did not necessarily represent the functional similarity of the bacteria and acquiring genomic data of geographical strains was essential to predict bacterial traits more accurately. Moreover, UMAP plots reflecting additional local structural variation (Fig. 2C, n_neighbors = 5) placed the strains G006c13, G009c41 and G010c01, G011c23 at distant positions, suggesting that obtaining strain-resolved genomes from each host is necessary to understand these differences.

### Acquisition of cSAGs from human gut bacteria using scALA

The strain-specific LR-SAGs were assembled using scALA. For nine *A. hadrus* strains, LR-SAGs with one-digit contig numbers were obtained, and for two *A. hadrus* strains, a single closed genome sequence of 3.12 Mbp (*A. hadrus* G009c41) and 3.30 Mbp (*A. hadrus* G011c23) was constructed without manual scaffolding. (Table S5).

To evaluate the assembly accuracy of *A. hadrus* LR-SAG constructed from SAGs containing chimeric sequences, we tested the alignment of LR-SAGs against the known complete genome of *A. hadrus* strain BPB5. The alignment results demonstrated that only the strain G012c17 LR-SAG was homologous to the strain BPB5 genome (Fig. S4). The other strains had a large inversion in the 1.7–2.2 Mb region of the BPB5 genome; however, the possibility of misassembly was exceptionally low because this structural variation was common to all eight LR-SAGs. Massive inversions of over 500 kbp are challenging to detect in SR-SAGs because they mainly consist of sequence fragments of tens of kbp or less. Finally, closed cSAGs of each strain (nine, four, and three genomes for *A. hadrus, R. gnavus*, and *A. rectalis*, respectively) were constructed by polishing and gap-filling between contigs of LR-SAG with scSR of the same samples and were used for subsequent analyses (Table 1).

**Table 1:** Circular single-cell amplified genomes (cSAGs) of *A. hadrus, A. rectalis*, and *R. gnavus*

| Species | Strain | Genome size (Mb) | GC (%) | Completeness (%) | Redundancy (%) | tRNA types | 16S rRNA | CDS |
| --- | --- | --- | --- | --- | --- | --- | --- | --- |
| *A. hadrus* | G004c06 | 3,251,112 | 36.79 | 95.64 | 2.52 | 19 | 6 | 3,112 |
| *A. hadrus* | G006c13 | 2,988,610 | 36.99 | 97.65 | 1.34 | 20 | 7 | 2,987 |
| *A. hadrus* | G009c41 | 3,098,900 | 37.06 | 98.99 | 2.68 | 20 | 6 | 2,966 |
| *A. hadrus* | G010c01 | 3,222,401 | 37.18 | 95.13 | 2.01 | 20 | 6 | 3,053 |
| *A. hadrus* | G011c23 | 3,312,423 | 37.18 | 99.33 | 2.01 | 20 | 5 | 3,141 |
| *A. hadrus* | G012c40 | 3,390,800 | 37.05 | 97.99 | 2.01 | 20 | 7 | 3,292 |
| *A. hadrus* | G012c17 | 3,214,912 | 37.21 | 99.33 | 2.68 | 20 | 6 | 3,057 |
| *A. hadrus* | G013c12 | 3,362,201 | 36.99 | 98.32 | 2.68 | 18 | 4 | 3,335 |
| *A. hadrus* | G015c54 | 3,259,135 | 36.93 | 94.21 | 2.01 | 20 | 6 | 3,237 |
| *R. gnavus* | G007c17 | 3,592,770 | 42.74 | 97.72 | 2.44 | 20 | 4 | 3,671 |
| *R. gnavus* | G007c21 | 3,276,103 | 42.65 | 90.62 | 6.14 | 19 | 5 | 3,224 |
| *R. gnavus* | G008c02 | 3,344,851 | 42.79 | 99.32 | 0 | 20 | 5 | 3,198 |
| *R. gnavus* | G010c06 | 3,418,459 | 43.07 | 96.37 | 0 | 20 | 4 | 3,338 |
| *A. rectalis* | G004c08 | 2,910,410 | 41.79 | 96.98 | 0 | 20 | 5 | 2,704 |
| *A. rectalis* | G011c14 | 3,718,229 | 41.63 | 98.55 | 0 | 20 | 5 | 3,599 |
| *A. rectalis* | G016c02 | 3,590,394 | 41.13 | 95.89 | 1.93 | 20 | 5 | 3,362 |

Furthermore, we analyzed the closed genome set of three bacterial species, consisting of cSAGs obtained in this study and one reference circular genome for each species (CP012098, NC012781, and CP043051). From the results of the pan-genome analysis using the closed genome set, we generated a genome plot of the three species using genome alignment based on homologous gene positions (Fig. 3A). Multiple strain-specific sequence regions of up to 100 kbp or more were identified throughout the genome. The pan-genome analysis of each bacterial species revealed that *A. hadrus* strains shared among cohabitants (G010c01-G011c23 and G012c40-G013c12-G015c54) had five-fold fewer unique genes than those of host-specific *A. hadrus* strains (Fig. 3B), which highlighted that gut bacteria were shared among cohabitants. Functional annotation of gene sequences obtained from each strain genome using Kyoto Encyclopedia of Genes and Genomes (KEGG)^20^ and Virus Orthologous Groups (VOG)^21^ confirmed that phage-like genes were significantly concentrated in strain-specific sequence regions. The rate of VOG-annotated genes in accessory or unique genes was higher than that in core genes (p-value, accessory vs core: 0.042, unique vs core; 0.0081) (Fig. 3C). Guanine-cytosine (GC) contents of the core genome showed a single peak, whereas GC contents of strain-specific regions containing viral sequences showed a more widely spread distribution with multiple weak peaks (Fig. 3D).

**Figure 3:**
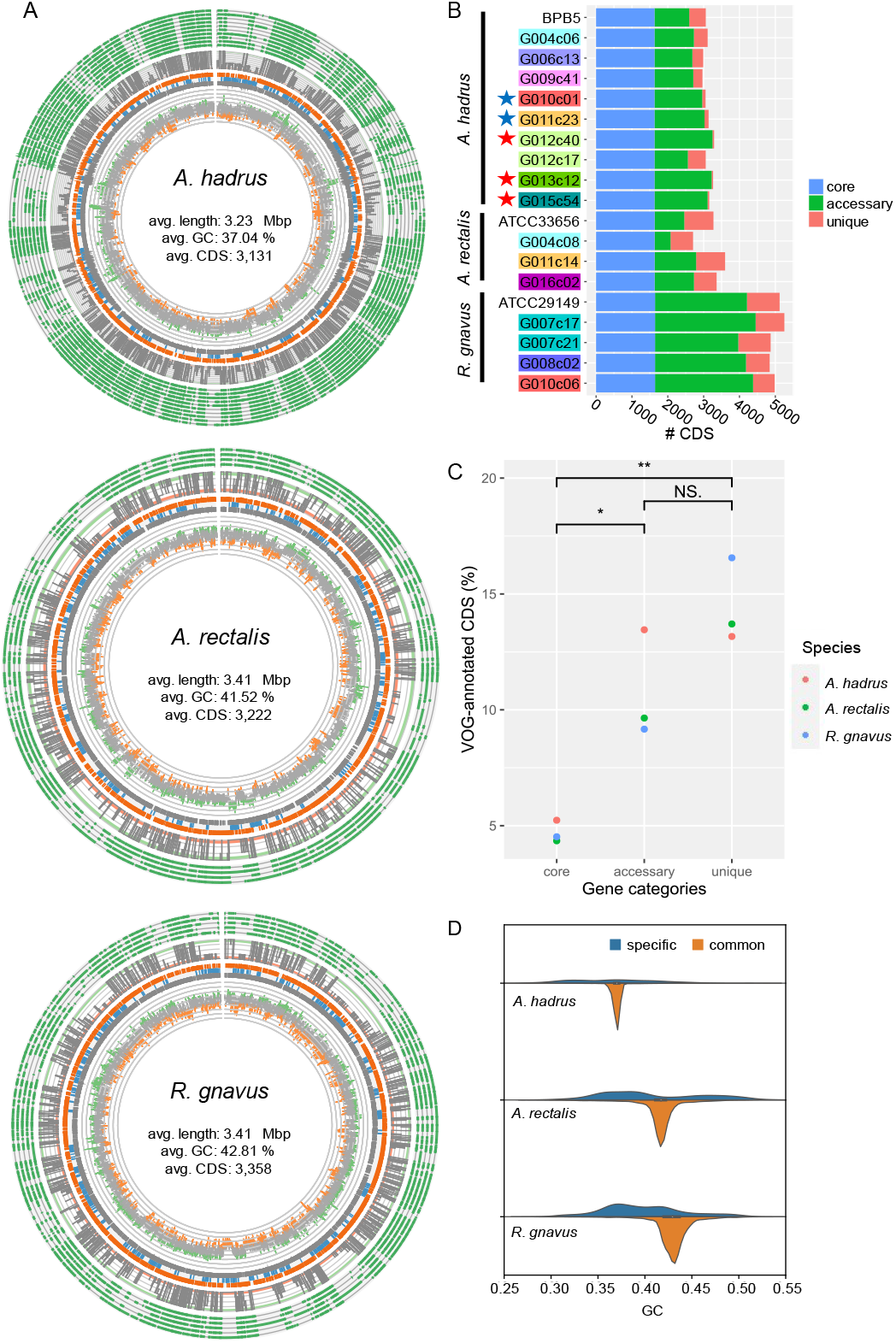
Comparison of the structures of gut bacterial circular single-cell amplified genomes (cSAGs). **A)** Genome maps of *A. hadrus, A. rectalis*, and *R. gnavus*. Track 1 (outside) tiles indicate the presence (green) or absence of gene cluster in each strain genome. Track 2 is a line plot indicating the number of genomes, including the gene cluster (green ribbon: core gene, orange ribbon: unique gene). Track 3 plots the gene cluster annotations (orange: Kyoto Encyclopedia of Genes and Genomes (KEGG)-annotated clusters, blue: COG-annotated clusters, gray: unannotated clusters). Track 4 is a line plot of GC contents (green: high-GC region, orange: low-GC region). **B)** Number of homologous gene clusters in cSAGs and reference genomes. The number of coding sequences (CDS) in core orthologous genes (OGs) (blue), accessory OGs (green), and unique OGs (red) is shown per cSAG. Highlighted labels indicate the hosts, and colored stars indicate closely related genomes obtained from the cohabitants. **C)** Accumulation of virus orthologous groups (VOG)-annotated CDS in strain-specific accessory or unique categories. **D)** GC content distributions of three bacterial genomes divided into strain-specific and strain-common sequence regions.

### Metabolic analysis of strain-specific structural variations observed in cSAGs

Metabolic analysis using obtained cSAGs was conducted to examine the manner in which structural variations affect bacterial traits. The screening of functional gene modules showed strain-specific differences in the presence or absence of gene modules associated with CAZy (Carbohydrate-Active enZymes), clustered regularly interspaced short palindromic repeats (CRISPR), and certain functions (Fig. 4A). CAZy profile indicated that some strains had the metabolic capacity of more diverse carbohydrates compared with that of other strains of the same bacterial species. For example, amorphous cellulose, beta-mannan, xylans, and xyloglucan metabolic capacities were detected only in the strain G004c06 of *A. hadrus* cSAG set, and only strains G006c13, G012c40, G013c12, G015c54 showed rhamnose metabolic capacity.

**Figure 4:**
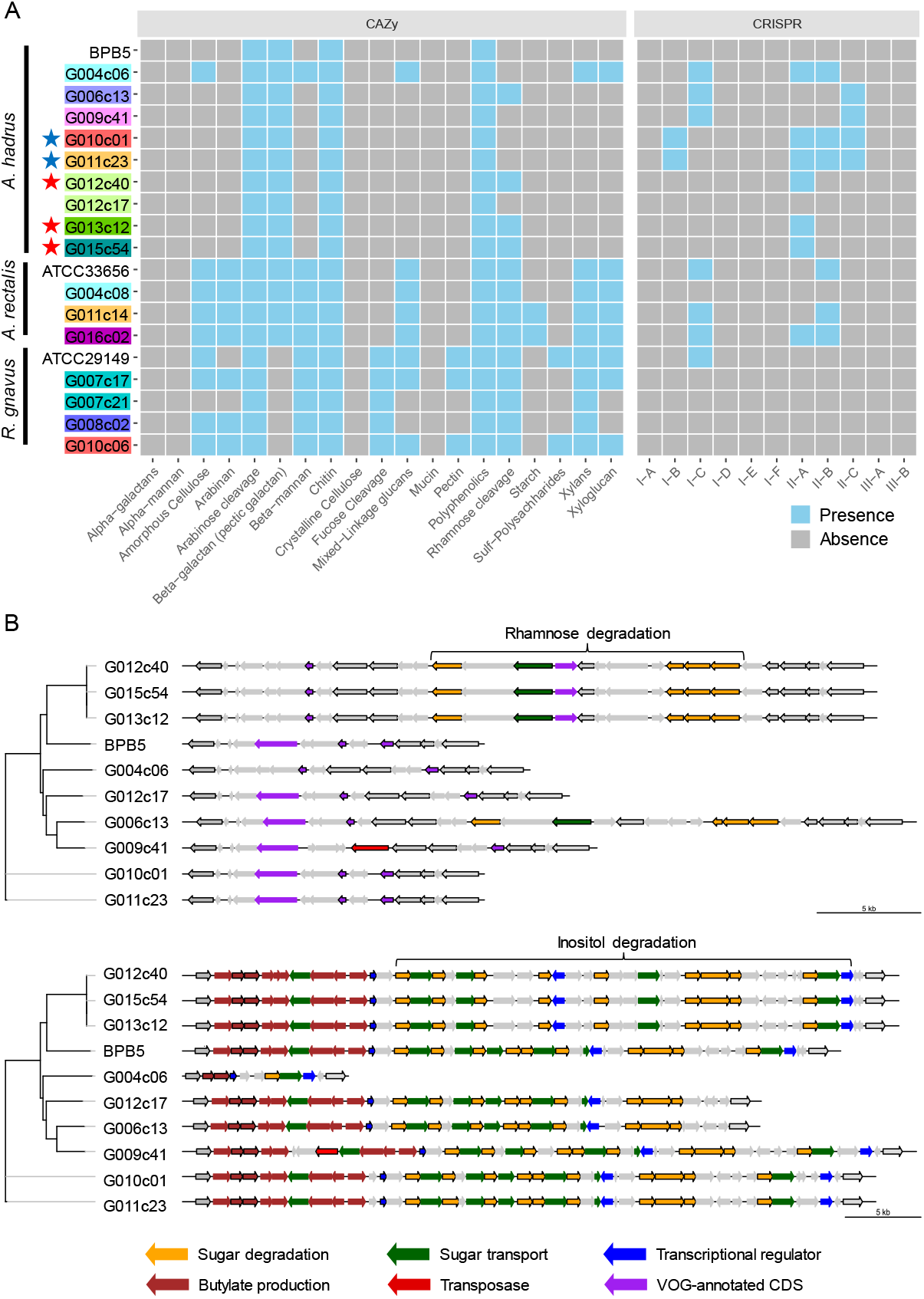
Metabolic analysis of gut bacterial circular single-cell amplified genomes (cSAGs). **A)** Heatmap showing various carbohydrate-active enzymes (CAZy) or CRISPR-Cas systems in the genome of each strain. Highlighted labels indicate the host, and colored stars indicate closely related genomes obtained from the cohabitants. **B)** Structural variation of polysaccharide metabolic pathways in *A. hadrus* genomes. Gene clusters in the same regions are aligned. The left phylogram was inferred from the present pattern of orthogroups in whole genomes.

The presence of the gene module related to rhamnose metabolism was visualized using *A. hadrus* cSAGs (Fig. 4B). The inositol metabolic system is a gut bacterial metabolic system highlighted in a cohort study using shotgun metagenomic data of the human gut microbiota^19^, suggesting the presence or absence of inositol metabolism gene module in the *A. hadrus* genome was correlated with host body weight and metabolic disease risks. In the above mentioned study, metagenomic reads were mapped to a database of bacterial genomes to detect variable and deletion structural variants based on differences in read depth; all inositol metabolism genes in *A. hadrus* were deleted in approximately 40 % of their metagenomic data sets. In contrast, the results of our study showed more diverse structural variations, such as gene rearrangements and partial deletions of functional genes, as well as the presence or absence of the entire inositol metabolic system in *A. hadrus* (Fig. 4B). In addition, such as the rhamnose metabolic system analysis, inserted gene sequences could not be detected using mapping-based evaluation owing to the loss in the reference *A. hadrus* BPB5 genome.

The number of CRISPR arrays in each genome varied: 2–7, 2–4, and 0–1 loci in the *A. hadrus, A. rectalis*, and *R. gnavus* genomes, respectively (Fig. 4A). *A. hadrus* genomes were classified into five types based on possession pattern of CRISPR-Cas systems, and transposase genes near the CRISPR-Cas systems indicated that these systems were transferred horizontally. The same types of CRISPR-Cas systems were located on the same regions of *A. hadrus* genomes, suggesting that each CRISPR-Cas system was inherited from the same ancestor strain acquired in the CRISPR-Cas system. Interestingly, CRISPR array sequences exhibited high strain specificity.

### Tracing genome structure dynamics of *A. hadrus*

Nine months after collecting fecal samples containing *A. hadrus* G011c23, a fecal sample was collected again from the same host, and seven SR-SAGs of *A. hadrus* were obtained using SAG-gel platform. The newly obtained SR-SAGs of *A. hadrus* strain G001c10 demonstrated > 99.5 % ANI with G011c23 SR-SAGs; therefore, genomic structural variations that occurred over time were investigated. CSR-SAGs constructed from *A. hadrus* G001c10 SR-SAGs were compared with *A. hadrus* G011c23. The G001c10 CSR-SAG had a 50-kbp highly diverse genomic region, including a 20-kbp deletion (Fig. 5A) and 12-kbp insertion (Fig. 5B), at the position corresponding to 2.3 Mbp of the G011c23 cSAG. Although traces of repeat sequences at both ends of the insertion region were observed, indicating the region was a prophage sequence (Fig. 5B), low homology of repeat sequences suggested that the detected structural variation occurred prior to the last nine months. This highly diverse region in the strain G001c10 was similar to the sequence structure in the strain G010c01 genome obtained from a close relative of the strain G011c23 host (Fig. 5A). Moreover, the sequence structures shared only between G011c23 and G001c10 were identified; thus, the three strains, G010c01, G011c23, and G001c10, were considered distinct. Ortholog (OG) analysis revealed that the strain G001c10 had significantly fewer specific OGs (28) than those of the strain G010c01 or G011c23 (169 and 226, respectively), indicating that the strain G001c10 genome had intermediate genome homology with the strains G010c01 and G011c23 (Fig. 5C). Structural differences were also detected in individual SAGs, suggesting that a single strain was dominant in each sample or at each sampling time. The genome sequences of the three strains, including existing marker sequences such as 16S rDNA genes, were very similar, barring relevant structural variation regions, suggesting that the variation in the dominant *A. hadrus* strain was newly detected using comprehensive scLR sequencing.

**Figure 5:**
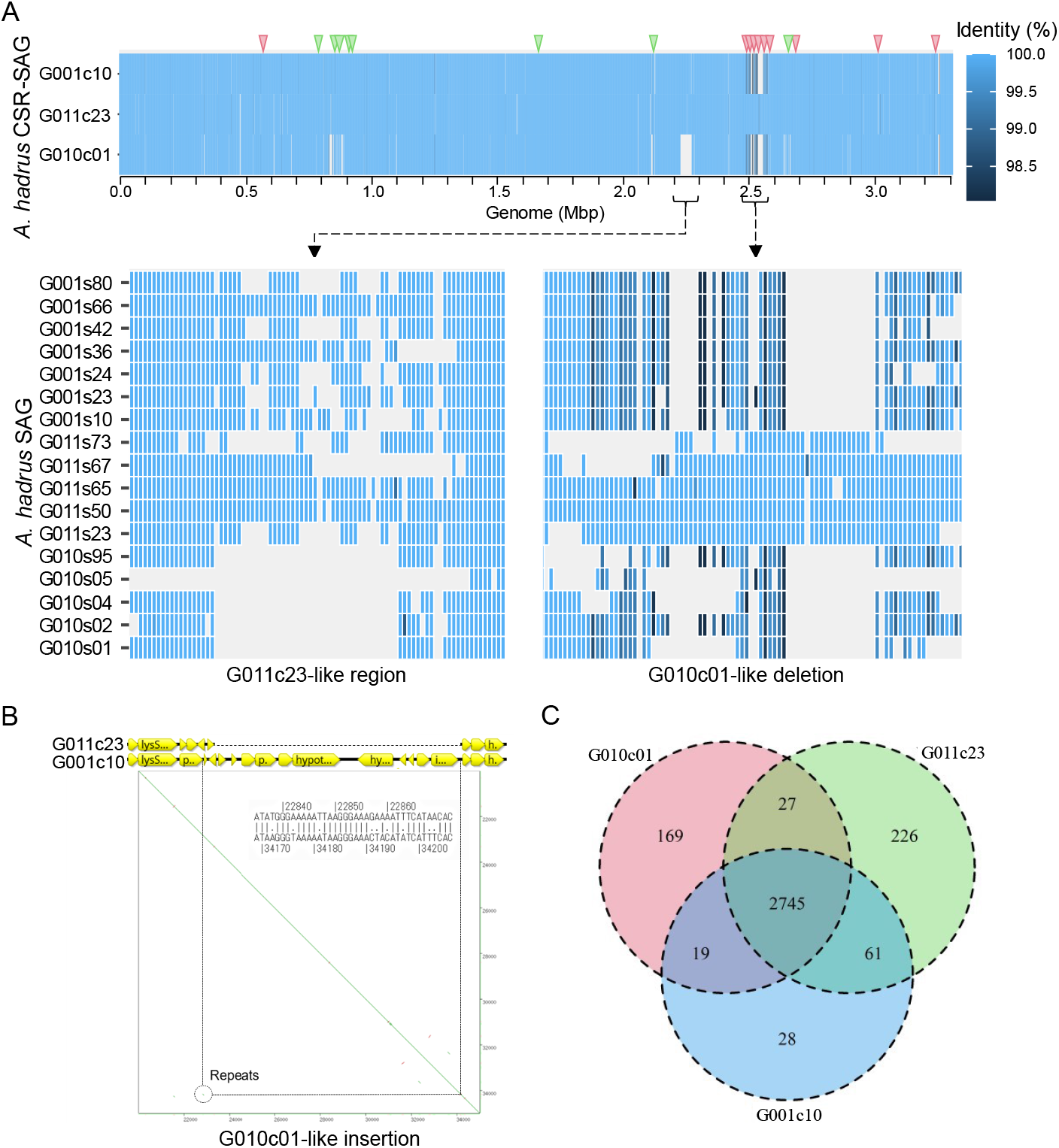
Inter-strain genome structure variation in *A. hadrus* strain SAGs obtained from the same host or a cohabitant at different time points. **A)** Heatmap showing nucleotide identity of each strain CSR-SAG with the circular single-cell amplified genomes (cSAGs) of *A. hadrus* G01c23. G011c23 and G001c10 derived from the same host, whereas strain G010c01 was derived from the cohabitant. Triangles at the top of the heatmap indicate SNPs detected upon comparing the three *A. hadrus* strains (red: G001c10-G010c01 common nucleotides, green: G001c10-G011c23 common nucleotides). Detailed view shows raw SR-SAG mapping results, indicating G010c01-like deletion in the G001c10 genome compared with the G011c23 genome. **B)** Gene map and dot plot of G010c01-like insertion region in the G001c10 genome compared with the G011c23 genome. **C)** Benn diagram of homologous gene clusters in genomes of the three *A. hadrus* strains.

## Discussion

Obtaining accurate and complete target bacterial genomes from multiple strains is important for studying the characteristics of uncultured environmental bacterial strains. Although advances in long-read sequencing and associated analysis technologies have made it possible to assemble circular bacterial genomes from cultured strains or metagenomes, obtaining complete bacterial genomes by combining long-read sequencing with uncultured bacterial SAGs was challenging. One reason for this difficulty was that the single-cell genome was not uniformly amplified during the whole genome amplification reaction; thus, low-frequency amplified genomic regions were missed from the assembled draft genomes. In this study, we used scALA and obtained circular human gut bacterial SAGs. In particular, we constructed the SAGs by iteration of existing genome assembly algorithms and mapping-based debiasing using scLR sequencing.

Multiple SAGs derived from the same bacterial strains that exhibit high levels of genome quality are required to construct uncultured bacterial cSAGs using scALA. Because the genome completeness of individual SAGs rarely reached 100 % and most SAGs had a genome completeness of 40–60 %, long-read sequencing of pooled samples of multiple SAGs is required to cover whole genomic information. Genome completeness and amplification bias exhibited by each SAG in short-read sequencing results should be considered when pooling SAGs and determining the mixing ratio. However, acquiring multiple qualified specific-strain SAGs is challenging owing to low genome quality, high running costs, and the complexity of experimental operations in conventional single-cell sequencing. In this study, we used cost-effective and high-throughput single-cell sequencing SAG-gel platform^11^ to easily obtain hundreds of SR-SAGs and multiple LR-SAGs of specific gut bacterial strains with the required quality for scALA.

Comparative genomics of *A. hadrus*, *A. rectalis*, and *R. gnavus* cSAGs revealed strain-unique structural variations, whereas they showed high homology in the aligned genomic regions. The strain-specific structural variations could be challenging to validate with conventional metagenomic approaches, including 16S rRNA gene amplicon sequencing and mapping short-read metagenomic sequencing reads to known reference genomes. Structural variations can be critical in determining the phenotype of individual bacterial strains, including the presence or absence of the gene sets such as sugar metabolism system, CRISPR-Cas system, or a synthetic flagellar module (Table S6). Moreover, our results indicated that analyzing CRISPR-Cas systems could reveal a more accurate evolution and distribution of *A. hadrus* than conventional marker gene analysis could. Metagenomic long-read sequencing used to construct circular bacterial genomes has been reported recently; however, the conventional accuracy of long-read sequences and assembly algorithms often render binning metagenomic sequences from the closely related bacterial species challenging^12^. Therefore, obtaining circular genomes of specifically targeted gut bacterial species using single-cell genome sequencing will likely be helpful in characterizing bacteria and detecting specific structural variations and gene sets in their genomes.

*A. hadrus*, whose genome was obtained in this study, is a notable host health-associated gut bacteria candidate^19^. Numerous studies on gut bacteria have been conducted, and the gut bacterial genome database is expanding^3^. These studies have also suggested the importance of investigating both representative genome sequences of specific species and geographically specific genome features. In this study, we confirmed that sequence similarity of *A. hadrus* genomes did not necessarily correlate with the presence of orthologous functional genes, whereas host geographical regionality appeared to be highly related to gene possession. In contrast, the *A. rectalis* genome highly correlated with sequence similarity, gene possession, and geographical location of the host, which is consistent with a large-scale comparative analysis with more than 1,300 *A. rectalis* genomes^22^ (Fig. S5). Our results suggested that the accumulation of type-strain genomes per geographical region was essential for the functional prediction of gut bacteria, and that suitable geographical resolution of strain classification depended on bacterial species.

The *A. hadrus* G001c10 genome contained a mixture of sequences that were in the genomes of two strains collected nine months earlier, the strain G011c23 from the same host and strain G010c01 from the cohabitant. The position of structural variations and SNPs indicated that the G001c10 genome had a partially chimeric genome consisting of the G010c01 and G011c23 genomes (Fig. 5A). Therefore, the emergence of strain G001c10 might be caused by homologous recombination of the G010c01 and G011c23 genomes. Additionally, *A. rectalis* G004c08 and *R. gnavus* G008c02 genomes were obtained after nine months; however, their structures hardly changed (data not shown). These results indicated that structural variation in the *A. hadrus* genome occurred at a high frequency compared with that in other gut bacteria, resulting in a loss of correlation between sequence homology and gene patterns.

Single-cell genome sequencing generally involves two main challenges arising from whole-genome amplification: chimeric sequences^23^ and amplification bias^24^. Thus, we developed scALA for *de novo* assembly of scLR sequences into circular SAG with addressing these problems. ScLR sequencing with scALA could be used to obtain complete circular genomes of uncultured bacteria in humans, soil, and marine and other polar environments to assess genomic structural variations. Structural variation such as inversions of promoter regions and gene sequences lead to changes in gene expression and phenotypes^25^. Moreover, obtaining complete genomes facilitates finding full-length prophage sequences and elucidates bacteria-phage interactions^26^. Our results suggest that strain-specific genomic regions formed owing to multiple host-specific phage infections that occurred in the past. Moreover, our study highlights the importance of the continuous accumulation of bacterial strain-resolved genomes to identify gene module alterations within species and understand their association with bacterial metabolic traits. We expanded the possibilities of uncultured strain genome comparison to identify novel genes and structural variations, which is rendered challenging using known reference genome-based metagenomic analyses^19^. The method we developed could identify unknown bacterial genomes using the closed genomes of single uncultured bacterial cells and enhance our understanding of intra-species diversities.

## Methods

### Experimental design and sample collection

The participants signed written informed consent, and the Ethics Review Committee of the Waseda University has approved the study (No. 2018-323). All methods used were conducted in accordance with the guidelines and regulations outlined by the ethics committee. The participants collected fresh feces into 15-mL vials containing 3 mL of guanidine thiocyanate (GuSCN) solution (TechnoSuruga Laboratory Co., Ltd.), and the samples were stored at 25 °C for a maximum of two days prior to single-cell encapsulation in droplets.

*E. coli* strain K-12 (ATCC 10798, genome size: 4.6 Mbp) was used for model analysis of cultured cells. *E. coli* K-12 cells were pre-cultured in Luria-Bertani medium for 16 h and collected using centrifugation. The collected cells were washed thrice with ultraviolet-treated Dulbecco’s Phosphate-Buffered Saline (DPBS, Thermo Fisher Scientific).

### Single-cell genome sequencing using SAG-gel platform

Single-cell genome amplification was performed using the SAG-gel platform, according to our previous report^11,27^. For gut bacteria analysis, after homogenization of human feces in phosphate buffered saline (PBS) or GuSCN solution (500 μL), the supernatant was recovered by centrifugation at 2,000 ×g for 30 s, followed by filtration through a 35-μm nylon mesh and centrifugation at 8,000 ×g for 5 min. The resulting cell pellets were resuspended in PBS and centrifuged twice at 8,000 ×g for 5 min.

Prior to single-cell encapsulation, cell suspensions were adjusted to 0.1 cells/droplets in 1.5 % agarose in PBS to prevent encapsulation of multiple cells in single droplets. Using an On-chip Droplet Generator (On-chip Biotechnologies Co., Ltd.), single bacterial cells were encapsulated in droplets floating in a carrier oil, 2 % Pico-Surf 1 in Novec 7500 (Dolomite Microfluidics) and were collected in a 1.5-mL tube, which was chilled on ice for 15 min to form the gel matrix. Following solidification, the collected droplets were broken with 1H,1H,2H,2H-perfluoro-1-octanol (Sigma-Aldrich) to collect the capsules. The gel capsules were washed with 500 μL of acetone (FUJIFILM Wako Pure Chemical Corporation), and the solution was mixed vigorously and centrifuged at 2,000 ×g for 10 seconds. The acetone supernatant was removed, 500 μL of isopropanol (FUJIFILM Wako Pure Chemical Corporation) was added, and the solution was once again mixed vigorously and centrifuged at 2,000 ×g for 10 seconds. The isopropanol supernatant was then removed, and the gel capsules were washed thrice with 500 μL of DPBS.

Thereafter, individual cells in beads were lysed by submerging the gel beads in lysis solutions: first solution, 50 U/μL Ready-Lyse Lysozyme Solution (Lucigen), 2 U/mL Zymolyase (Zymo Research Corporation), 22 U/mL lysostaphin (Sigma-Aldrich), and 250 U/mL mutanolysin (Sigma-Aldrich) in DPBS at 37 °C overnight; second solution, 0.5 mg/mL achromopeptidase (FUJIFILM Wako Pure Chemical Corporation) in DPBS at 37 °C for 6-8 h; and third solution, 1 mg/mL Proteinase K (Promega Corporation) in 0.5 % SDS in DPBS at 40 °C overnight. The lysis steps were skipped in *E. coli* experiment. At each reagent replacement step, the gel beads were washed thrice with DPBS and then resuspended in the next solution. After the lysis, the gel beads were washed with DPBS five times, and the supernatant was removed. Thereafter, the beads were suspended in Buffer D2 (denaturation buffer) and subjected to MDA reaction using a REPLI-g Single Cell Kit (QIAGEN).

After MDA at 30 °C for 2 h and 65 °C for 3 min, the gel beads were washed thrice with 500 μL of DPBS. The beads were then stained with 1× SYBR Green (Thermo Fisher Scientific) in DPBS. After confirming DNA amplification based on the presence of green fluorescence in the gel, fluorescence-positive beads were sorted into 0.8 μL of DPBS in 96-well plates using a BD FACSMelody cell sorter (BD Bioscience) equipped with a 488-nm excitation laser. After droplet sorting, 96-well plates proceeded through the second-round MDA or were stored at −30 °C.

Following gel bead collection in 96-well plates, second-round MDA was performed using the REPLI-g Single Cell Kit. Buffer D2 (0.6 μL) was added to each well and incubated at 65 °C for 10 min. Thereafter, 8.6 μL of MDA mixture was added and incubated at 30 °C for 120 min. MDA reaction was terminated by heating at 65 °C for 3 min. After second-round amplification, master library plates of SAGs were prepared. For quality control, SAG aliquots were transferred to replica plates, which were used for DNA yield quantification using a Qubit dsDNA High Sensitivity Assay Kit (Thermo Fisher Scientific).

### Short-read sequencing

For sequencing analysis, scSR libraries were prepared from the second-round MDA product using QIAseq FX DNA Library Kit (QIAGEN). The ligation adapters were modified to TruSeq–Compatible Full-length Adapters UDI (Integrated DNA Technologies, Inc.). Each SAG library was sequenced using an Illumina HiSeq 2× 150 bp configuration (Macrogen).

Furthermore, scSRs quality-controlled with fastp 0.20.0^28^ (option: -q 25 -r -x) were assembled to SR-SAG *de novo* using SPAdes 3.14.0 (options for SAG: --sc −careful −disable-rr −disable-gzip-output -t 4 -m 32), and contigs of less than 1,000 bp were excluded from subsequent analyses^23^. SR-SAGs with the completeness of > 10 % and contamination of < 10 % were selected using CheckM v1.1.2^29^. ANI was calculated for the selected SR-SAGs using FastANI 1.3^30^. The homology of common single-copy marker genes obtained using CheckM taxonomy workflow (option: -nt −tab_table -t 16 domain Bacteria) was calculated using blastn 2.9.0+ with default options. From individual host datasets, SAGs with ANI of > 95 %, single-copy marker gene homology of > 99 %, and tetra-nucleotide frequencies correlation of > 90 % were classified in the same strain group. The SR-SAGs in strain groups were integrated into CSR-SAGs using ccSAG for chimera removal and co-assembly^9^. Unless otherwise stated, the analysis tools were run with default parameters.

### Long-read sequencing and standard *de novo* assembly

We prepared scLR libraries from individual *E. coli* SAGs using the Rapid Sequencing Kit (Oxford Nanopore Technologies) and sequenced them with Flow Cell R9.4 using a GridION (Oxford Nanopore Technologies). *E. coli* scLRs (300 Mbp) were obtained from five *E. coli* second-round MDA products and then integrated into a single file.

We selected second-round MDA products of fecal bacteria from 96-well plates based on strains identified using CSR-SAGs and mixed them (2–8 SAGs per strain) for scLR sequencing using a nanopore sequencer. We used Miniasm 0.3^15^, Canu 1.9^16^, and Flye 2.7^14^ for the assembly of scLRs into LR-SAGs. LR-SAG quality was assessed using QUAST 5.0^31^ and CheckM. The alignment results of the draft genome and reference *E. coli* genome were visualized using D-GENIES 1.2.0^32^.

### LR-SAG assembly using scALA

We removed low-quality scLRs of < 1,000 bp or with an average quality score of < 10 after removing the first 75 bases from the input scLRs using Nanofilt^33^. Thereafter, we constructed reference contigs for scLR debiasing. All assemblies in scALA were performed by chimeric sequence trimming using Canu 1.9^16^ and subsequent *de novo* assembly of scLRs using Flye 2.7^14^. The input scLRs were mapped against reference contigs using minimap2 2.17^15^. Biased sequences were identified according to mapped depth, and the excess number of the input scLRs was removed to be 50× sequence depth. By re-assembling using the bias-reduced LR-SAGs, we constructed the second cycle reference contigs that cover a more comprehensive range of genomic regions than do the first reference contigs. We repeated the assembly and debiasing processes four times. Consensus sequences were then constructed based on the alignment of multiple reference contigs obtained during the bias-reduction-assembly cycle. Before and after the alignment, the sequences were corrected using Pilon 1.22^34^ and scSRs obtained from the same MDA product. After sequence polishing, we obtained strain-specific LR-SAGs. Closed cSAGs of *A. hadrus, R. gnavus*, and *A. rectalis* were constructed by polishing and gap-filling LR-SAG contigs with scSR of the same samples.

### Comparative genome analysis

We performed pan-genome analysis using Roary^35^ with reference genomes in the NCBI genome database. Clustering of strains was performed using UMAP analysis based on the presence of homologous gene groups^36^. Distance matrices of the genomes were generated using the Manhattan method and visualized using the R package umap 0.2.7^37^.

Genome alignment results of LR-SAGs were visualized using D-1.2.0^32^ or Circos 0.69^38^ for bacterial genomic structural variation analysis. For phylogenetic analysis, Multiple Alignment using Fast Fourier Transform-based alignment was performed for concatenated sequences of homologous genes detected in the target genomes, and phylogenetic trees generated with the maximum likelihood method using RAxML-NG 0.9.0^39^ were illustrated using iTOL 6.4.3^40^. Furthermore, KEGG annotation with Kofamscan^20^ and VOG annotation with VIBRANT 1.2.1^21^ were conducted against coding sequences (CDSs) extracted using Prokka 1.14.5^41^ annotation and CDS positions were compared with structural variation sites. The t-test of VOG gene possession was conducted with R. Gene modules were illustrated using genoplotR^42^. Pathway analysis and CRISPR detection were conducted using DRAM 1.0.6^43^ and CRISPRDetect 2.4^44^, respectively.

## Declarations

### Ethics approval and consent to participate

Studies with human participants were approved by the ethics review committee at Waseda University (No. 2018-323). The participants provided written informed consent prior to sample collection.

### Consent for publication

Not applicable.

### Availability of data and material

Sequencing data have been deposited in the NCBI database under BioProject PRJNA818799 and PRJNA692334.

## Acknowledgments

The super-computing resource was provided by the Human Genome Center (University of Tokyo).

## Authors’ contributions

MK and MH conceived and designed the experiments. MK, YN, HT, and MH developed the long-read single-cell sequencing. MK, TS, and TY conducted the genomics experiments and collected the data. MK and KA conducted bioinformatics analysis of single-cell genomic data. MK and MH analyzed all data and wrote the original manuscript. All authors reviewed and approved the final manuscript.

## Supplementary files

**Figure S1** Coverage of *E. coli* genome assembled from debiased sequence reads. Read depth in each *E. coli* genomic region of scLR before (top) and after sequence debiasing at depth = 50 (bottom). The red lines indicate sequence regions that are not included in the draft genome obtained from the same scLR assembly method using Canu and Flye.

**Figure S2** Alignment of intermediate draft assembled in multistep debiasing. Vertical gray lines show the edge of contigs aligned to the reference *E. coli* genome (strain K-12, substrain MG1655). Intermediate draft genomes constructed for debiasing of long reads has contig sequences that are interrupted at different positions from each other. The interrupted position (red arrow) of the intermediate draft assembled in the later cycle may be contiguous (green arrow) in the earlier intermediate draft.

**Figure S3** Network showing average nucleotide identity of *A. hadrus* SAGs. Rectangles are the SAGs, and the color is assigned to each host participant. SAGs with > 99.5% ANI is linked with purple line, and SAGs with > 98% ANI is linked with gray line.

**Figure S4** Alignment of CoSAG-LR draft genomes of *A. hadrus* species. Horizontal axis shows the position of the reference genome (GenBank ID: CP012098.1). There were 500-kbp large inversion at the 1.7-2.2 Mb region of the reference genome in the 8 out of 9 acquired LR-SAGs *A. hadrus* draft genomes.

**Figure S5** Phylogenetic analysis using *A. rectalis* genomes. Publicly available *A. rectalis* genomes obtained from different countries were used for comparison with our SAGs. **A)** a parSNP tree of *A. rectalis* based on single nucleotide variants (SNPs) of the core genes. Color of the labels shows the countries where the bacterial genome was acquired. The right color strip shows the genome group based on the UMAP analysis (**B**). The UMAP analysis of the *A. rectalis* genomes was implemented based on the presence of homologue gene groups. The point color shows the region where the bacterial sample was acquired, and the point shape shows the data type of the genome sequences.

**Table S1** scLR genome assembly using existing long-read assemblers

**Table S2** Genome assembly of *Escherichia coli* with scALA

**Table S3** Host information of gut bacteria

**Table S4** CoSAG of three gut microbial species

**Table S5** LR-SAGs of three gut microbial species

**Table S6** Functional annotation with DRAM please refer to the supplementary document.

