## Supplementary Information for "Revealing within-species diversity in uncultured human gut bacteria with single-cell long-read sequencing"

### Contents

**Supplementary Figure S1-S5** 2

**Supplementary Table S1-S5** 7

#### Supplementary Figure S1-S5

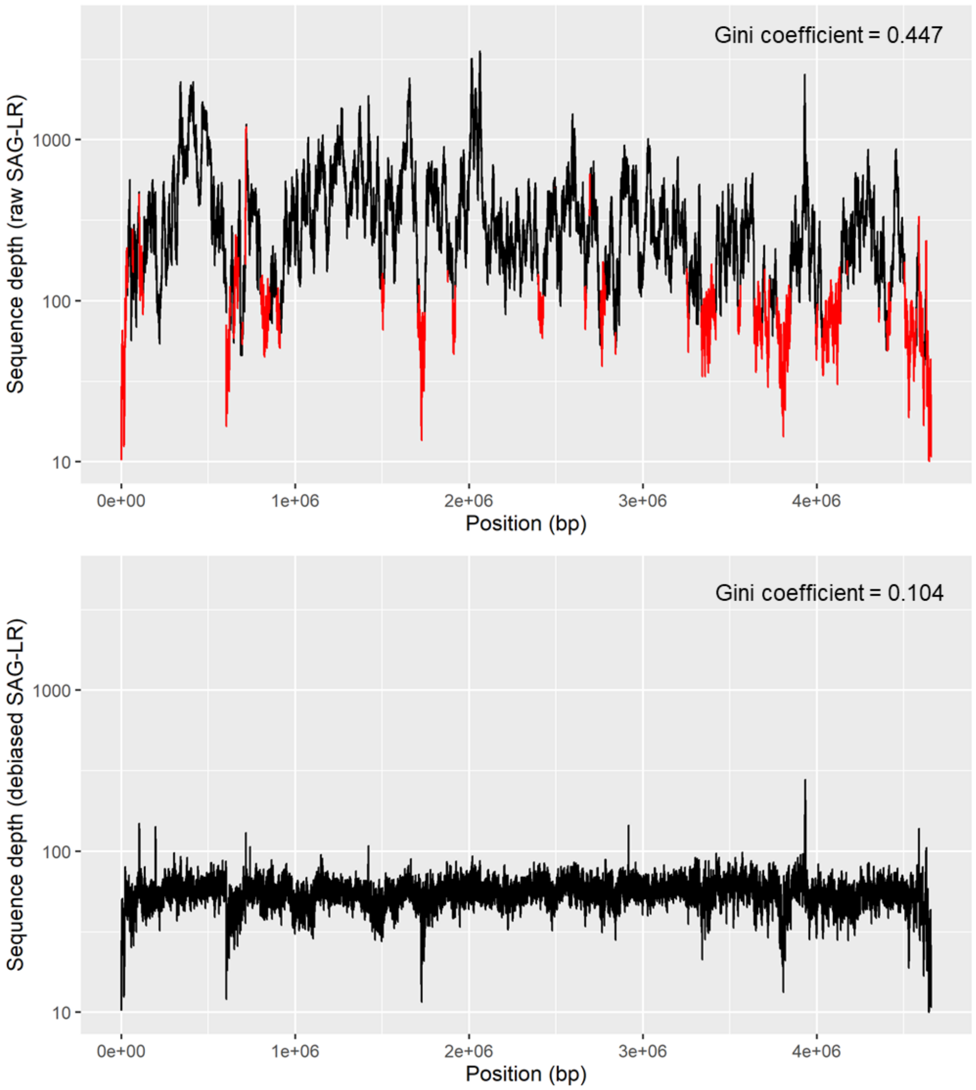


**Figure S1:** Coverage of *E. coli* genome assembled from debiased sequence reads. Read depth in each *E. coli* genomic region of scLR before (top) and after sequence debiasing at depth = 50 (bottom). The red lines indicate sequence regions that are not included in the draft genome obtained from the same scLR assembly method using Canu and Flye.

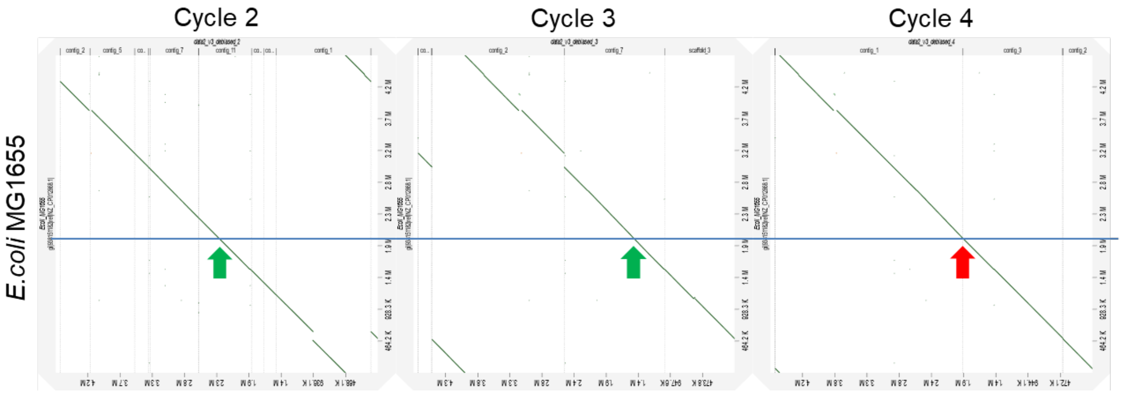


**Figure S2**: Alignment of intermediate draft assembled in multistep debiasing. Vertical gray lines show the edge of contigs aligned to the reference *E. coli* genome (strain K-12, substrain MG1655). Intermediate draft genomes constructed for debiasing of long reads has contig sequences that are interrupted at different positions from each other. The interrupted position (red arrow) of the intermediate draft assembled in the later cycle may be contiguous (green arrow) in the earlier intermediate draft.

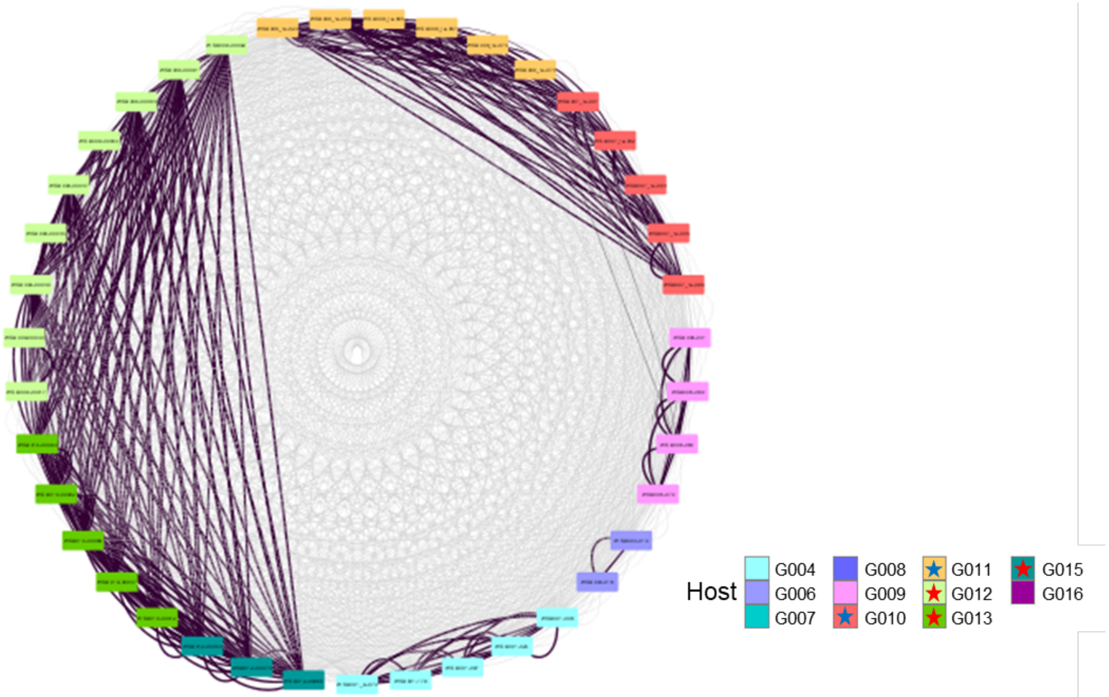


**Figure S3**: Network showing average nucleotide identity of *A. hadrus* SAGs. Rectangles are the SAGs, and the color is assigned to each host participant. SAGs with > 99.5% ANI is linked with purple line, and SAGs with > 98% ANI is linked with gray line. The colored stars in the host legends indicate cohabitants.

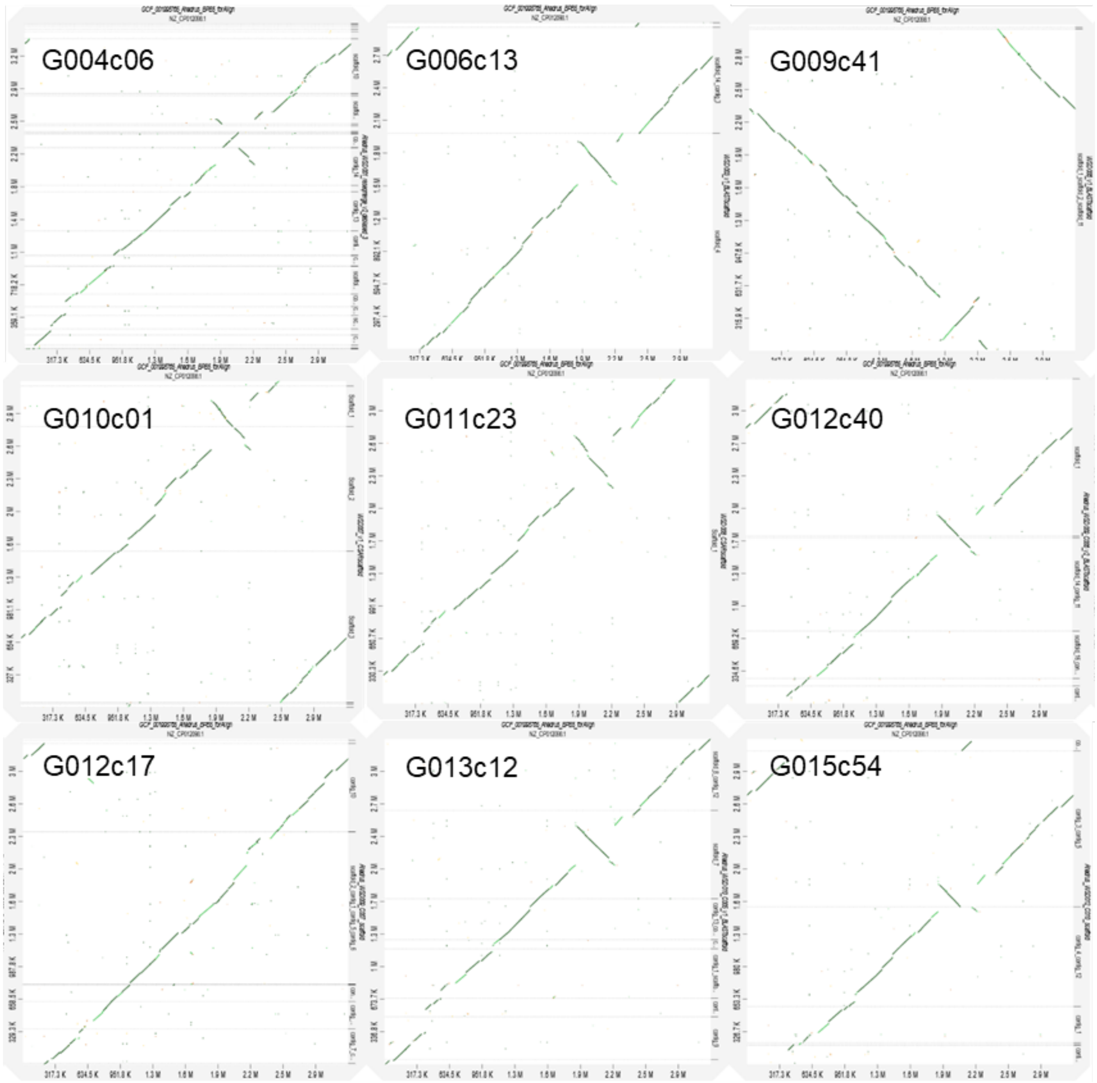


**Figure S4:** Alignment of CoSAG-LR draft genomes of *A. hadrus* species. Horizontal axis shows the position of the reference genome (GenBank ID: CP012098.1). There were 500-kbp large inversion at the 1.7-2.2 Mb region of the reference genome in the 8 out of 9 acquired LR-SAGs *A. hadrus* draft genomes.

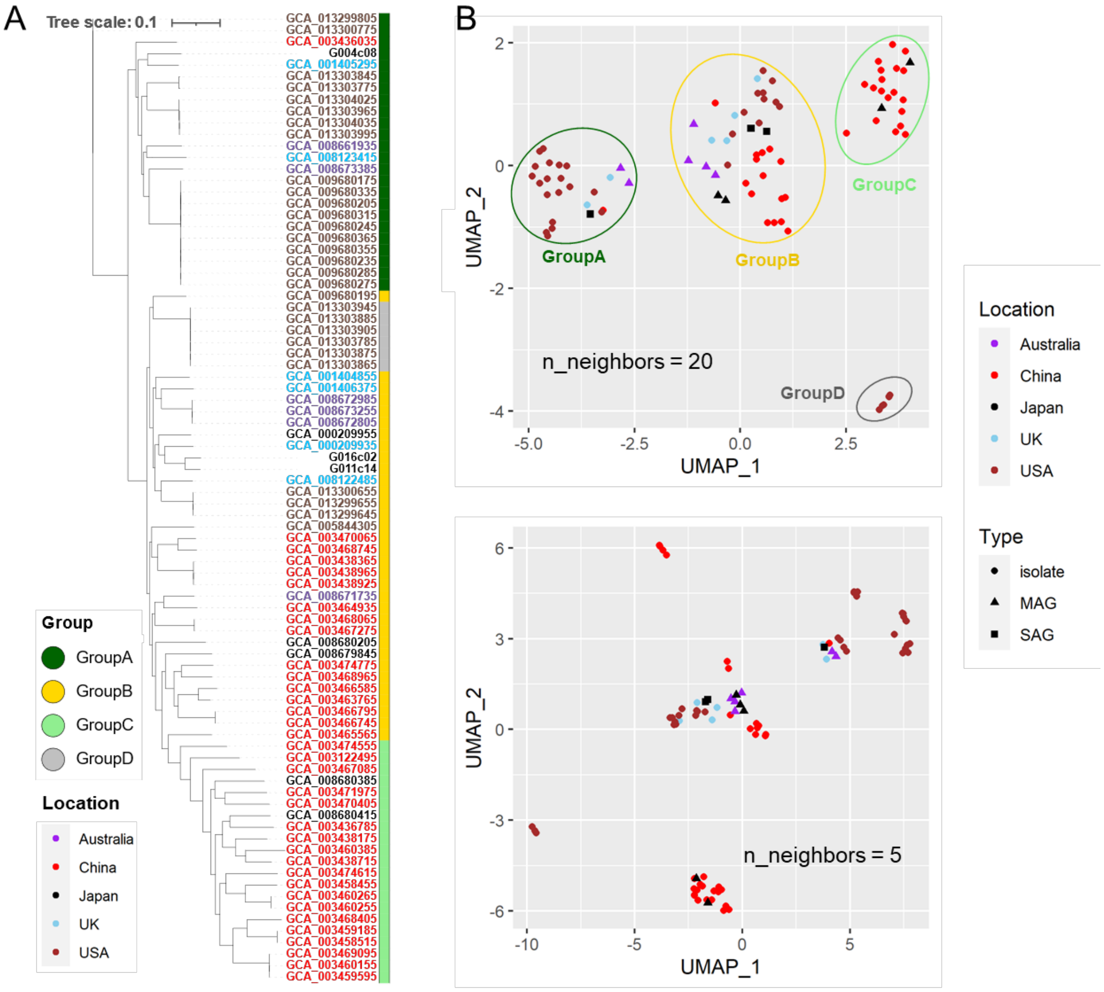


**Figure S5:** Phylogenetic analysis using *A. rectalis* genomes. Publicly available *A. rectalis* genomes obtained from different countries were used for comparison with our SAGs. **A)** a parSNP tree of *A. rectalis* based on single nucleotide polymorphisms (SNPs) of the core genes. Color of the labels shows the countries where the bacterial genome was acquired. The right color strip shows the genome group based on the UMAP analysis (**B**). The UMAP analysis of the *A. rectalis* genomes was implemented based on the presence of homologue gene groups. The point color shows the region where the bacterial sample was acquired, and the point shape shows the data type of the genome sequences.

#### Supplementary Table S1-S5

For

**Table S6:** Functional annotation with DRAM,

please refer to the supplementary document.

**Table S1:** scLR genome assembly using existing long-read assemblers

| Assembler | Input LR | # contigs | Total length (bp) | Largest contig (bp) | NG50 (bp) | Genome fraction (%) | Duplication ratio |
|---:|---:|---:|---:|---:|---:|---:|---:|
| Flye | Raw | 3 | 91,461 | 41,356 | 36,686 | 1.531 | 1.356 |
| Miniasm | Raw | 25 | 1,424,444 | 155,249 | 60,710 | 1.174 | 1.001 |
| Canu | Raw | 61 | 4,223,998 | 579,808 | 216,678 | 87.501 | 1.039 |
| Flye | Chimera removed | 39 | 3,877,368 | 585,307 | 197,181 | 81.671 | 1.003 |
| Miniasm | Chimera removed | 47 | 3,355,167 | 468,842 | 123,007 | 70.543 | 1.001 |

**Table S2:** Genome assembly of *Escherichia coli* with scALA

| Assembly | # contigs (>= 5 kbp) | Total length | Largest contig (bp) | NG50 (bp) | Genome fraction (%) | Duplication ratio |
|---:|---:|---:|---:|---:|---:|---:|
| Trial-a Cycle-0 | 39 | 3,877,368 | 585,307 | 130,901 | 81.4 | 1.003 |
| Trial-a Cycle-1 | 13 | 4,700,839 | 1,307,120 | 649,334 | 98.528 | 1.002 |
| Trial-a Cycle-2 | 3 | 4,737,661 | 3,179,672 | 3,179,672 | 98.857 | 1.005 |
| Trial-a Cycle-3 | 2 | 4,717,848 | 4,650,553 | 4,650,553 | 98.894 | 1.001 |
| Trial-a Cycle-4 | 2 | 4,725,667 | 4,658,362 | 4,658,362 | 99.028 | 1.002 |
| Trial-a Scaffold | 2 | 4,725,667 | 4,658,362 | 4,658,362 | 99.028 | 1.002 |
| Trial-b Cycle-0 | 43 | 5,154,123 | 522,358 | 275,957 | 82.989 | 1.315 |
| Trial-b Cycle-1 | 40 | 4,539,545 | 560,233 | 215,879 | 95.232 | 1.004 |
| Trial-b Cycle-2 | 8 | 4,710,942 | 2,391,184 | 2,391,184 | 98.764 | 1.001 |
| Trial-b Cycle-3 | 3 | 4,720,075 | 3,158,259 | 3,158,259 | 98.926 | 1.002 |
| Trial-b Cycle-4 | 5 | 4,730,521 | 2,388,484 | 2,388,484 | 98.7 | 1.006 |
| Trial-b Scaffold | 2 | 4,701,452 | 4,634,168 | 4,634,168 | 98.095 | 1.006 |
| Trial-c Cycle-0 | 39 | 4,370,605 | 483,518 | 232,914 | 81.566 | 1.131 |
| Trial-c Cycle-1 | 28 | 4,818,271 | 751,821 | 291,066 | 97.434 | 1.044 |
| Trial-c Cycle-2 | 11 | 4,676,808 | 1,380,328 | 706,662 | 98.231 | 1.001 |
| Trial-c Cycle-3 | 6 | 4,734,508 | 1,948,980 | 1,479,831 | 98.735 | 1.007 |
| Trial-c Cycle-4 | 5 | 4,717,134 | 2,752,764 | 2,752,764 | 98.829 | 1.002 |
| Trial-c Scaffold | 2 | 4,716,838 | 4,649,538 | 4,649,538 | 98.872 | 1.001 |

**Table S3:** Host information of gut bacteria

| Series | Host | Sampling Date | Cohabitant | Sex | Age |
|---:|---:|---:|---:|---:|---:|
| G002 | WSD008 | 2020/05/26 | - | male | 36 |
| G004 | WSD001 | 2019/09/02 | - | male | 33 |
| G006 | WSD003 | 2019/09/25 | - | male | 44 |
| G007 | WSD004 | 2019/09/25 | - | male | 35 |
| G008 | WSD005 | 2019/09/25 | - | male | 28 |
| G009 | WSD006 | 2019/09/29 | A | male | 2.3 |
| G010 | WSD007 | 2019/09/29 | A | female | 30 |
| G011 | WSD008 | 2019/09/29 | A | male | 35 |
| G012 | WSD009 | 2020/01/13 | B | female | 11 |
| G013 | WSD010 | 2020/01/14 | B | male | 45 |
| G015 | WSD012 | 2020/01/14 | B | female | 43 |
| G016 | WSD013 | 2019/09/30 | - | male | 45 |

**Table S4:** CoSAG of three gut microbial species

| Species | Strain | # contigs (>= 500 bp) | Largest contig (bp) | Total length (bp) | GC (%) | N50 (bp) | Completeness (%) | Redundancy (%) |
|---:|---:|---:|---:|---:|---:|---:|---:|---:|
| *A. hadrus* | G004c06 | 269 | 96,526 | 3,101,238 | 36.87 | 30,117 | 98.32 | 1.12 |
| *A. hadrus* | G006c13 | 239 | 92,221 | 2,807,309 | 36.82 | 27,918 | 95.41 | 1.34 |
| *A. hadrus* | G009c41 | 224 | 111,663 | 3,110,996 | 37.33 | 39,574 | 99.33 | 2.68 |
| *A. hadrus* | G010c01 | 236 | 122,807 | 3,084,635 | 37.17 | 31,056 | 97.99 | 2.75 |
| *A. hadrus* | G011c23 | 194 | 122,807 | 3,116,373 | 37.05 | 35,247 | 99.33 | 2.01 |
| *A. hadrus* | G012c40 | 176 | 112,747 | 3,257,254 | 36.96 | 37,613 | 99.33 | 2.68 |
| *A. hadrus* | G012c17 | 444 | 109,772 | 3,796,778 | 38.26 | 30,437 | 99.27 | 11.74 |
| *A. hadrus* | G013c12 | 184 | 202,461 | 3,253,384 | 36.94 | 37,229 | 99.33 | 2.35 |
| *A. hadrus* | G015c54 | 250 | 130,894 | 3,207,170 | 36.94 | 33,172 | 96.48 | 2.18 |
| *R. gnavus* | G007c17 | 232 | 160,919 | 3,155,029 | 42.61 | 36,663 | 96.28 | 1.17 |
| *R. gnavus* | G007c21 | 438 | 94,286 | 3,010,460 | 42.59 | 18,519 | 80.45 | 1.46 |
| *R. gnavus* | G008c02 | 143 | 216,621 | 3,159,004 | 42.82 | 55,519 | 98.39 | 0 |
| *R. gnavus* | G010c06 | 274 | 206,908 | 3,450,926 | 43.1 | 38,161 | 98.83 | 1.85 |
| *A. rectalis* | G004c08 | 133 | 228,950 | 2,779,296 | 41.74 | 45,920 | 96.7 | 0.02 |
| *A. rectalis* | G011c14 | 243 | 113,875 | 3,428,283 | 41.62 | 42,584 | 98.83 | 0 |
| *A. rectalis* | G016c02 | 285 | 111,486 | 3,402,239 | 41.26 | 30,688 | 98.93 | 1.13 |

**Table S5:** LR-SAGs of three gut microbial species

| Species | Strain | # reads | read size (Mbp) | average read length (bp) | maximum read length (bp) | # contigs (>= 10 kbp) | # contigs (>= 50 kbp) | Largest contig (bp) | Total length (bp) | GC (%) | N50 (bp) |
|---:|---:|---:|---:|---:|---:|---:|---:|---:|---:|---:|---:|
| *A. hadrus* | G004c06 | 64,636 | 283 | 4,374.80 | 73,500 | 25 | 16 | 599,753 | 3,591,047 | 37.23 | 305,351 |
| *A. hadrus* | G006c13 | 462,575 | 1,222 | 2,642.30 | 96,827 | 3 | 2 | 1,963,467 | 2,973,587 | 37.02 | 1,963,467 |
| *A. hadrus* | G009c41 | 401,797 | 1,286 | 3,201.20 | 93,722 | 3 | 1 | 3,115,941 | 3,158,589 | 37.22 | 3,115,941 |
| *A. hadrus* | G010c01 | 242,903 | 821 | 3,380.40 | 95,017 | 6 | 4 | 1,515,008 | 3,270,179 | 37.17 | 1,251,086 |
| *A. hadrus* | G011c23 | 297,923 | 928 | 3,115.90 | 85,644 | 1 | 1 | 3,303,338 | 3,303,338 | 37.24 | 3,303,338 |
| *A. hadrus* | G012c40 | 263,765 | 860 | 3,261.90 | 97,800 | 6 | 5 | 1,620,223 | 3,345,933 | 36.97 | 955,999 |
| *A. hadrus* | G012c17 | 229,570 | 916 | 3,991.90 | 80,514 | 9 | 7 | 1,537,379 | 3,290,168 | 37.23 | 889,286 |
| *A. hadrus* | G013c12 | 227,123 | 900 | 3,521.30 | 84,116 | 8 | 8 | 916,679 | 3,368,464 | 37.14 | 514,282 |
| *A. hadrus* | G015c54 | 229,570 | 1,188 | 3,503.10 | 90,511 | 6 | 5 | 1,566,289 | 3,259,027 | 37.05 | 1,001,773 |
| *R. gnavus* | G007c17 | 381,715 | 1,208 | 3,163.40 | 98,412 | 19 | 10 | 906,164 | 3,892,163 | 42.51 | 393,456 |
| *R. gnavus* | G007c21 | 221,508 | 815 | 3,679.60 | 80,846 | 19 | 12 | 545,618 | 3,355,870 | 42.63 | 387,639 |
| *R. gnavus* | G008c02 | 288,344 | 971 | 3,366.20 | 73,841 | 6 | 1 | 3,369,411 | 3,546,440 | 42.88 | 3,369,411 |
| *R. gnavus* | G010c06 | 263,362 | 1,028 | 3,902.10 | 79,719 | 14 | 11 | 748,237 | 3,462,859 | 43.05 | 588,945 |
| *A. rectalis* | G004c08 | 224,517 | 819 | 3,650.10 | 79,298 | 4 | 3 | 2,418,663 | 2,945,069 | 41.78 | 2,418,663 |
| *A. rectalis* | G011c14 | 222,980 | 868 | 3,891.30 | 84,039 | 1 | 1 | 3,774,748 | 3,780,922 | 41.58 | 3,774,748 |
| *A. rectalis* | G016c02 | 310,082 | 1,081 | 3,485.70 | 81,028 | 12 | 8 | 1,147,138 | 3,626,467 | 41.14 | 758,463 |
